# Epigenetic Regulation of Inflammatory NF-κB Target Genes by IFN-γ via IRF1

**DOI:** 10.1101/2024.11.26.625557

**Authors:** Bikash Mishra, Claire Wingert, Mahesh Bachu, Lionel B. Ivashkiv

## Abstract

The regulation of inflammatory gene expression involves complex interactions between transcription factors (TFs), signaling pathways and epigenetic chromatin-mediated mechanisms. This study investigated mechanisms by which by IFN-γ-mediated priming augments TLR-induced expression of NF-κB target genes in primary human monocytes. IFN-γ priming enhanced the expression of signature inflammatory genes such as *IL6*, *TNF*, *IL1B*, and *CXCL10* when monocytes were exposed to various TLR agonists. RNA-seq analysis identified genes synergistically activated by IFN-γ and LPS, which were enriched in inflammatory pathways. Similar synergistic activation was observed with the TLR1/2 agonist PAM3CYS, suggesting a shared regulatory mechanism. ATAC-seq analysis revealed that TLR ligands induce IRF1 TF activity independently of IFN-γ. JAK1/2 inhibitor (iJAK) treatment reduced IRF1 expression and protein levels, especially in IFN-γ-treated monocytes, but not in LPS-stimulated monocytes, suggesting LPS-induced IRF1 may compensate for loss of IFN-γ-induced IRF1. We applied CRISPR-Cas9 to knock out IRF1 in primary human monocytes and found loss of IRF1 abrogates synergistic activation of key inflammatory genes, suggesting a pivotal role for IRF1. This genetic data was corroborated by IRF1 CUT&RUN data showing resistance of IRF1 binding to JAK inhibition under (IFN-γ + LPS) costimulated conditions, and co-occupancy of IRF1 binding sites by NF-κB. This study enhances our understanding of inflammatory gene regulation, highlighting IRF1 as a key player and a potential therapeutic target for inflammatory diseases.

## Introduction

Inflammation, a fundamental biological response, is essential for host defense against pathogens and repair of tissue injury ^1, 2^. However, when dysregulated, it can lead to the development of chronic inflammatory diseases such as rheumatoid arthritis (RA), inflammatory bowel disease (IBD), and systemic lupus erythematosus (SLE) ^2, 3, 4, 5^. Central to the inflammatory response are monocytes and macrophages, key innate immune cells that play pivotal roles in recognizing and responding to infectious agents and damaged tissues ^6, 7, 8^. These cells are activated by various stimuli, including cytokines and pathogen-associated molecular patterns (PAMPs), which trigger signaling cascades that ultimately lead to the induction of inflammatory genes ^1, 2, 8, 9, 10^.

One of the key mediators of inflammation is tumor necrosis factor (TNF), a proinflammatory cytokine produced primarily by activated monocytes/macrophages ^2, 11^. TNF plays a crucial role in the regulation of immune responses and is a major driver of inflammation in many chronic inflammatory diseases ^11, 12^. Toll-like receptors (TLRs) are another essential component of the innate immune system that recognize specific molecular patterns associated with pathogens ^1, 2, 10, 13, 14^. Ligands for TLRs, such as lipopolysaccharide (LPS) and PAM3CSK4 (PAM), can activate monocytes and macrophages, leading to the activation of NF-κB and production of inflammatory mediators.

Interferon-gamma (IFN-γ) is another key player in the inflammatory response, primarily produced by activated T cells and natural killer cells ^15, 16, 17^. IFN-γ is known to enhance the antimicrobial and inflammatory functions of macrophages and is involved in the regulation of immune responses ^18^. Priming of monocytes with IFN-γ has been shown to significantly enhance their responsiveness to secondary stimuli, a process that includes synergistic transcriptional activation of NF-κB target genes ^19, 20, 21, 22^. However, the precise molecular mechanisms underlying this synergistic activation of inflammatory genes in IFN-γ-primed monocytes are not fully understood.

Recent studies have highlighted the importance of epigenetic changes and chromatin remodeling in the regulation of gene expression during inflammation ^21, 22^. These changes can alter the accessibility of transcription factors (TFs) to their target genes, thereby influencing gene expression. TFs such as NF-κB, AP-1, and interferon regulatory factors (IRFs) play critical roles in the transcriptional regulation of inflammatory genes ^20, 21^. NF-κB is a key transcription factor involved in the regulation of inflammatory responses, and its activation can be triggered by a range of stimuli beyond just TLR ligands. Inflammatory stimuli such as cytokines (e.g., TNF-α), microbial products (e.g., bacterial, and viral components), and various stress signals can all activate the NF-κB pathway ^11^. Once activated, NF-κB translocates into the nucleus and regulates the expression of a wide array of genes involved in immune responses and inflammation. In the context of IFN-γ primed monocytes, the combination of IFN-γ priming and subsequent stimulation with other inflammatory signals could potentially lead to increased cooperation between STAT1 and IRFs downstream of IFN-γ, and NF-κB and AP-1 downstream of secondary stimuli, leading to enhanced activation of NF-κB target genes. However, the specific role of IRF1 in mediating the synergistic activation of inflammatory genes in IFN-γ-primed monocytes and its interaction with other TFs during TLR signaling pathways remains unclear.

Interferon regulatory factors (IRFs) are crucial in the immune response of monocytes, especially in cooperation with STATs to induce expression of interferon-stimulated genes (ISGs)^23^. The IRF family includes several members, each with unique roles. In addition to inducing ISGs, IRF1 has been implicated in expression of cytokines such as IL-12 and TNF-α during inflammation ^20, 22^. IRF3 and IRF7 are key in the antiviral response, promoting type I interferons (IFN-α and IFN-β) upon viral infection, typically activated through phosphorylation by kinases like TBK1 and IKKε, leading to their nuclear translocation and binding to interferon-stimulated response elements (ISREs) in target gene promoters ^23, 24^. IRF5 plays a significant role in activating pro-inflammatory cytokines via TLR signaling and can form complexes with transcription factors like NF-κB and AP-1 to enhance inflammatory gene transcription ^23, 24, 25, 26^. IRF8, often collaborating with IRF1, influences the differentiation and function of monocytes and macrophages and regulates various cytokines and chemokines ^27^. The interactions between different IRFs are complex, involving both cooperative and competitive mechanisms. For instance, IRF1 and IRF8 can amplify inflammatory responses, while IRF4 acts as a counter-regulatory factor, reducing excessive inflammation by inhibiting other IRFs and NF-κB ^23, 27^. This intricate IRF network ensures a balanced immune response, preventing overactivation that could lead to tissue damage and chronic inflammation. Understanding IRFs’ specific roles and interactions in monocytes can elucidate innate immune mechanisms and highlights potential therapeutic targets for inflammatory diseases.

In this study, we aimed to investigate the roles of TLR ligands in the activation of inflammatory genes in IFN-γ-primed primary human monocytes. Additionally, we wished to understand whether JAK inhibition can decrease induction of NF-κB target genes in IFN-γ- primed cells upon secondary challenge with inflammatory stimuli. We hypothesized that different TLR ligands may activate distinct sets of inflammatory genes and that IRF1 may play a crucial role in mediating the synergistic activation of these genes. To test this hypothesis, we employed a combination of techniques, including ATAC-seq to assess chromatin accessibility, RNA-seq to analyze gene expression, and CRISPR-Cas9-mediated gene editing to decrease IRF1 expression in primary human monocytes.

Interestingly, we found that IFN-γ priming alone was not able to elicit NF-κB target gene expression, but ATAC-seq data showed increased chromatin accessibility at these gene loci, suggesting IFN-γ primes chromatin for future activation. Notably, similar synergistic activation was observed in LPS and PAM challenged monocytes, highlighting a shared regulatory mechanism. Through ATAC-seq analysis, we observed that TLR ligands significantly induce IRF1 TF activity independently of IFN-γ. Using a JAK1/2 inhibitor (iJAK) baricitinib, we demonstrated that blocking IFN-γ signaling significantly reduces IRF1 levels and activity in IFN-γ-primed monocytes. Conversely, IFN-γ + LPS-challenged monocytes maintained IRF1 activity even with iJAK treatment.

To further investigate the role of IRF1, we employed CRISPR-Cas9 to knockout *IRF1* in primary human monocytes. The loss of IRF1 abrogated the synergistic activation of key inflammatory genes, such as TNF and ISG15, in the IFN-γ + LPS condition. Finally, CUT&RUN experiments revealed that IRF1 bound genomic regions that have binding sites for NF-κB transcription factors, suggesting cooperation in driving inflammatory responses. Overall, our findings provide strong genetic and epigenetic evidence that IRF1 plays a pivotal role in the synergistic activation of inflammatory genes in IFN-γ-primed human monocytes. The results underscore the importance of IRF1 in transcriptional memory and chromatin remodeling, contributing to the persistent high-level induction of NF-κB target genes. This study enhances our understanding of the molecular mechanisms underlying inflammatory gene regulation and identifies potential therapeutic targets for inflammatory diseases.

## Results

### RNAseq analysis reveals distinct and overlapping sets of genes superinduced by TLR4 and TLR2 signaling in IFN-γ-primed monocytes

We aimed to understand whether inflammatory stimuli activating the NF-κB pathway could super-activate these genes in IFN-γ-primed monocytes. To explore whether the regulatory effects of IFN-γ priming were confined to LPS-TLR4 mediated activation of inflammatory NF-κB target genes, we exposed IFN-γ-primed and unprimed monocytes to diverse NF-κB activators such as agonists of TLR1/2 (PAM), TLR4 (LPS), and TLR7/8 (VTX-2337, R848), and evaluated the mRNA levels of specific NF-κB target genes (Figure 1A). Like LPS challenged monocytes, IFN- γ-primed monocytes when stimulated with other TLR agonists synergistically activated signature inflammatory genes such as *IL6*, *TNF*, *IL1B* and ISGs like *CXCL10* (Figure 1B). Thus, super-activation of NF-κB target genes was induced by TLRs 4/7/8 that induce an autocrine IFN-β loop, and by TLR2 that does not.

**Figure 1:**
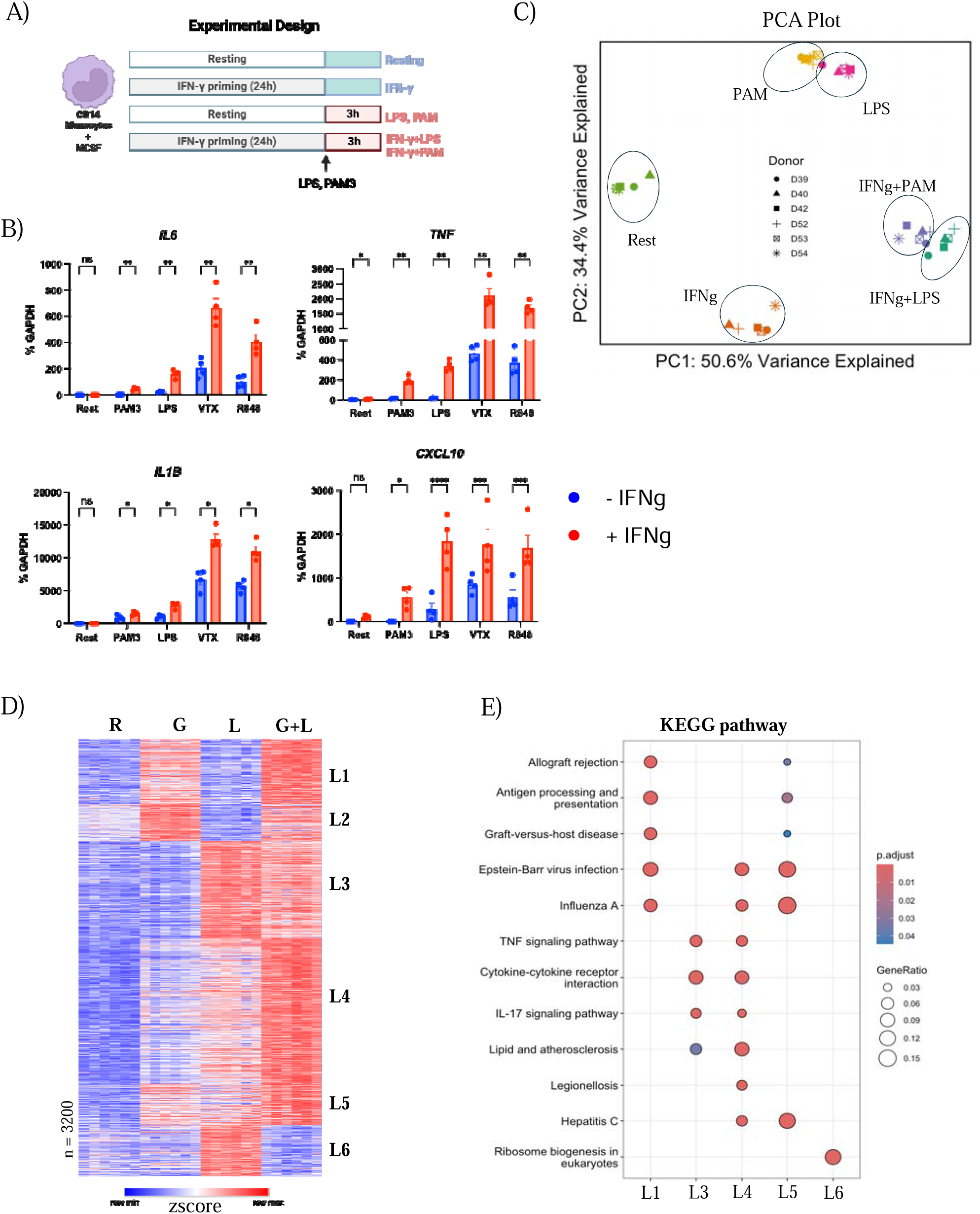
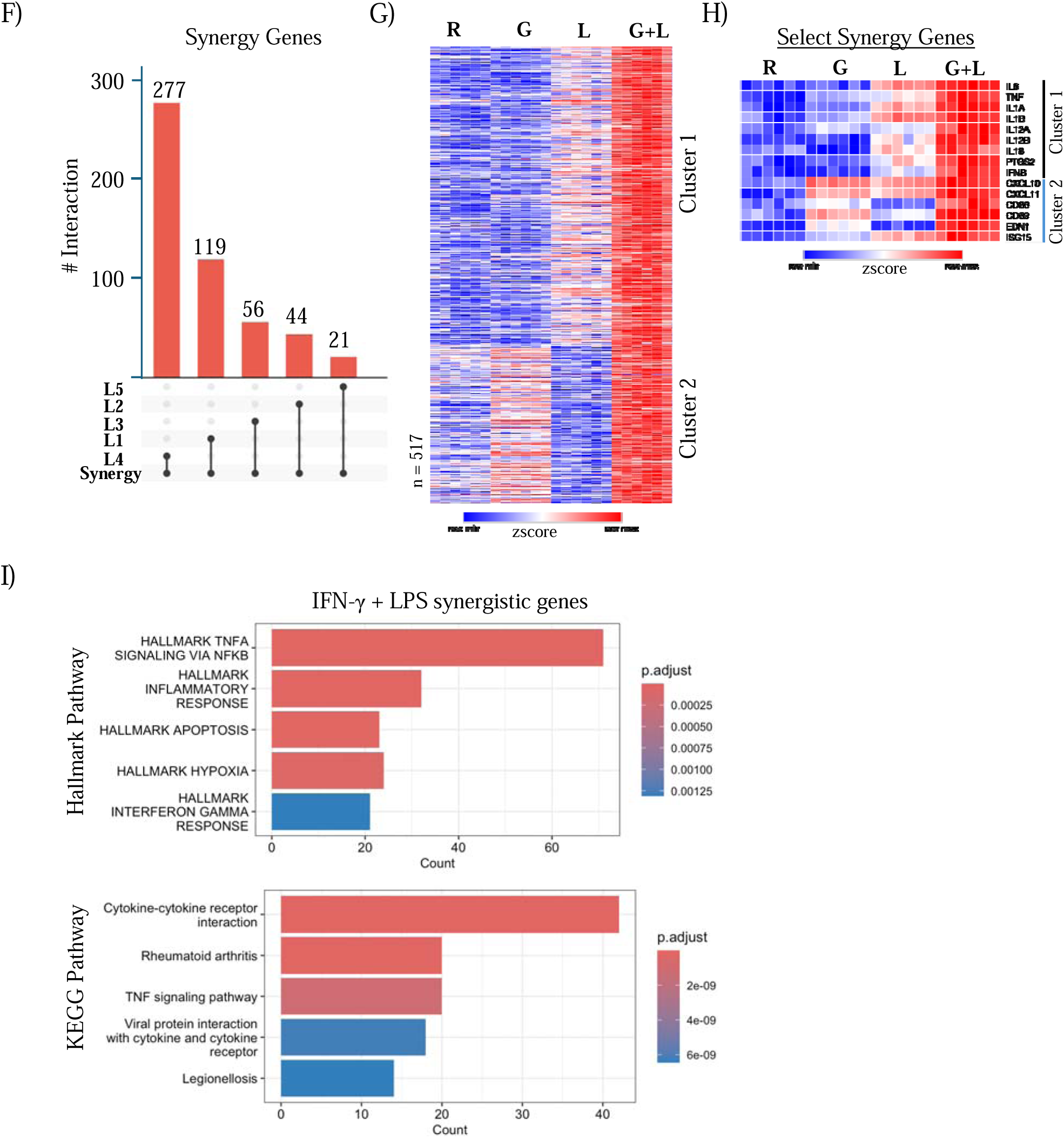
TLR ligands broadly induced synergistic activation of NF-kB target genes in IFN-γ primed monocytes. A) Experimental design. Human monocytes were cultured –/+ IFN-γ (100 U/mL) for 24 hours and then stimulated with various inflammatory ligands for 3 hours. B) mRNA of indicated genes was measured by qPCR and normalized relative to *GAPDH* mRNA in cells stimulated as indicated (dots correspond to independent donors; n = 4) C) PCA plot of RNA-seq data of IFN-γ-primed primary human monocytes challenged with indicated secondary stimuli. D) k-means clustering analysis (k=6) conducted on differentially upregulated genes in any pairwise comparison relative to resting control. R = Resting, G = IFN-γ, L = LPS, G + L = IFN-γ + LPS E) KEGG pathway enrichment analyses performed on clusters identified in Figure 2.1.D F) UpSet plot showing overlap between synergistically activated genes in IFN-γ-primed LPS-challenged monocytes and genes in individual k-means clusters from Figure 2.1.D G) Heatmap of synergistically activated gene in LPS model clustered by LPS vs IFN induced genes. H) Heatmap of subset of synergistically activated gene in LPS model from Figure 2.1.G I) Hallmark and KEGG pathway enrichment analyses performed on synergistically activated genes.

We performed RNA-seq analysis to identify the sets of genes that are synergistically activated across the genome when IFN-γ-primed monocytes are challenged with TLR1/2 agonist (PAM3CSK4 (PAM)), and TLR4 agonist (LPS) (Figure 1 C). Principal Component Analysis (PCA) showed treatment-specific effects, with distinct clustering for PAM, LPS, IFN-γ, and their combinations, indicating distinct transcriptional responses driven by each stimulus (Figure 1C). To characterize super-activated genes, we initially clustered genes based on their expression patterns and conducted pathway analysis. We focused first on LPS-challenged, IFN-γ-primed monocytes. In this group, genes in clusters labeled L1, L4, and L5 were super-induced under combined IFN-γ and LPS conditions, compared to their expression with either LPS or IFN-γ alone (Figure 1D). Conversely, clusters L2, L3, and L6 showed suppressed/reduced gene expression under the combined conditions. Pathway analysis revealed that super-induced clusters were strongly enriched for the inflammatory NF-κB pathway, IFN response genes, and processes downstream of IFN signaling (Figure 1E). Additionally, we utilized a previously published computational approach ^21^ to identify synergistically induced genes (greater than additive induction). Using this method, we identified 517 genes that were synergistically induced by IFN- γ + LPS. A substantial majority (417 out of 517, approximately 80%) of the synergistic genes identified through this computational approach were found within clusters L1, L4, and L5 (Figure 1F), indicating a robust overlap and reinforcing the validity of these genes as key targets in our study. The synergistic genes (n = 517) were categorized into two clusters based on their expression patterns (Figure 1G). Cluster one (n = 337) closely aligns with the expression profile of “signature” priming genes, which are minimally responsive to IFN-γ alone but significantly upregulated in primed cells challenged with LPS relative to resting cells challenged with LPS (Figure 1H, top panel). In contrast, cluster two (n = 180) genes are partially induced by IFN-γ alone and were then synergistically enhanced by the combined effect of IFN-γ priming and LPS challenge (Figure 1H, bottom panel). Pathway analysis of the core set of IFN-γ + LPS synergistic genes revealed a highly significant enrichment of pathways related to Rheumatoid arthritis and TNF-NF-κB signaling, with a less pronounced enrichment of IFN response pathways (Figure 1I). Our findings build on and corroborate previous reports ^19, 21, 22^ of IFN-γ + LPS synergistic genes, offering a comprehensive characterization of the full spectrum of synergy genes in human monocytes.

We then performed a similar RNA-seq analysis to determine if PAM-induced NF-κB target genes were super-activated in IFN-γ-primed monocytes, similar to their response to LPS. In the model with IFN-γ-primed and PAM-challenged monocytes (IFN-γ + PAM), clusters P1 and P3 displayed patterns similar to those observed in the LPS super-activated cluster, involving genes linked to the inflammatory NF-κB pathway, IFN response genes, and downstream IFN signaling processes (Figure 2 A-B). The similarity between pathways enriched suggests a common set of synergistically activated genes in the LPS and PAM models. We identified 400 genes that were synergistically induced by IFN-γ + PAM, with a substantial number (317 out of 400, or 79%) located in clusters P1 and P3 (Figure 2 C). These genes, like those in the LPS model, were significantly enriched for pathways associated with rheumatoid arthritis and TNF-NF-κB signaling (Figure 2 D).

**Figure 2:**
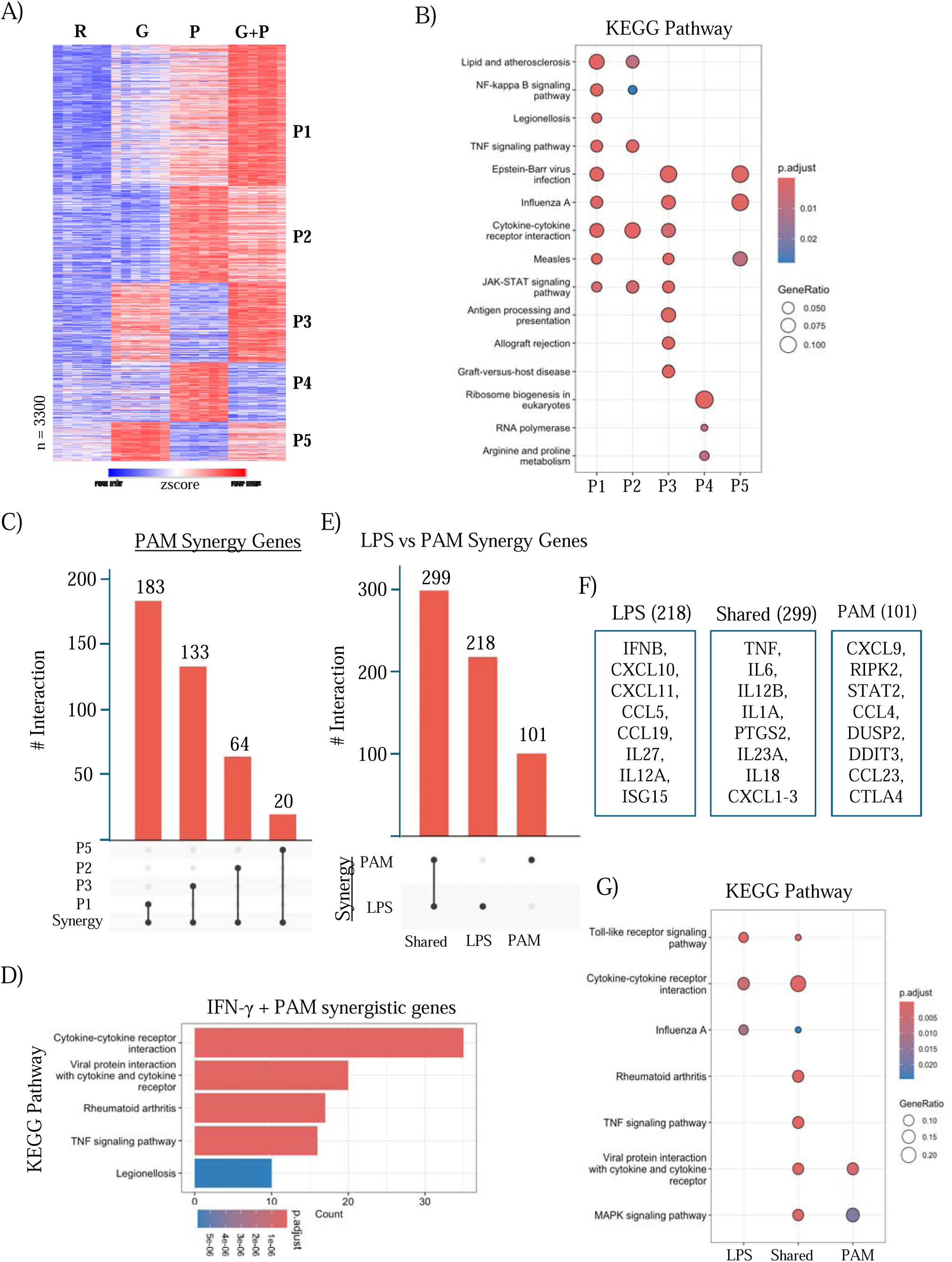
Overlap between TLRs induced synergistic activation of NF-κB Target genes in IFN-γ- primed and TLR-Challenged Monocytes. A) k-means clustering analysis (k=5) conducted on differentially upregulated genes in any pairwise comparison relative to resting control. R = Resting, G = IFN-γ, P = PAM, G + P = IFN-γ + PAM B) KEGG pathway enrichment analyses performed on clusters identified in Figure 2.2.A C) UpSet plot showing overlap between synergistically activated genes in IFN-γ-primed PAM-challenged monocytes and genes in individual k-means clusters from Figure 2.2.A D) KEGG pathway enrichment analyses performed on synergistically activated genes. E) UpSet plot showing overlap between synergistically activated genes in IFN-γ-primed PAM-challenged and LPS-challenged monocytes. F) Representative synergistically induced genes present in each group from Figure. 2.2.E.

A direct comparison of synergistically induced genes between the LPS and PAM models revealed that 299 (approximately 49%) of the 618 total synergy genes were common between both models, while 218 (35%) were uniquely synergistic in the LPS model, and 101 (16%) were uniquely synergistic in the PAM model (Figure 2 E). Notably, synergy genes activated in both the LPS and PAM models included “signature” priming genes, which are typically canonical NF-κB targets such as *TNF, IL6, IL12B, IL1A, PTGS2, IL23A,* and *IL18*. Genes uniquely synergistic in the LPS model included inflammatory cytokines and chemokines, *IFNB* and IL-12 family genes *IL12A* and *IL27*, and IFN response genes like *CXCL10, CXCL11, CCL5, CCL19* and *ISG15*; this pattern of superinduction only by LPS may be related to the LPS-induced IFN-β-mediated autocrine loop. Conversely, genes uniquely synergistic in the PAM model included immune mediator genes like *CXCL9, RIPK2, STAT2, CCL4, DUSP2/8/10/18, DDIT3*, and *CCL23* (Figure 2 F). Pathway analysis indicated an enrichment of TNF and rheumatoid arthritis pathway genes within the shared synergistic genes, implying that both LPS and PAM can synergistically activate core inflammatory genes implicated in inflammatory disorders such as rheumatoid arthritis (RA) (Figure 2 G).

### ATACseq analysis implicates IRFs in the IFN-γ-primed TLR response

We aimed to decipher the regulatory mechanisms leading to the synergistic activation of inflammatory genes following TLR signaling in IFN-γ-primed monocytes. To achieve this, we conducted ATAC-seq experiments to assess chromatin accessibility and performed motif enrichment analysis under ATAC-seq peaks to identify changes in transcription factor activity. To determine if IFN-γ priming increased chromatin accessibility at the regulatory regions of synergistic genes, we performed differential peak analysis and compared the differentially upregulated peaks (fold change ≥ 2, FDR ≤ 0.05) detected by ATAC-seq in the IFN-γ condition to the differentially upregulated genes detected by RNA-seq. This approach allowed us to correlate chromatin accessibility changes with transcriptional activation, shedding light on the epigenetic landscape and transcription factor dynamics driving the synergistic gene activation in response to combined IFN-γ and TLR signaling (Figure 3 A-B). We initially annotated IFN-γ-induced peaks to assign these peaks to genes and found that 7,932 peaks were associated with 2,825 genes. Next, we investigated whether increased chromatin accessibility corresponded with increased transcription. To test this, we utilized an upset plot analysis, comparing genes associated with IFN- γ-induced peaks from ATAC-seq data to IFN-γ-induced genes identified through RNA-seq. This analysis allows us to determine the overlap between chromatin accessibility changes and gene expression changes. We found that the majority of IFN-γ-induced accessible chromatin peaks did not correspond to IFN-γ-induced gene expression (2281 out of 3521, or 64.8%), indicating that not all regions of induced open chromatin result in gene activation (Figure 3 C). Figure 3D depicts representative gene tracks showing de novo ATAC peaks at *TNF* and *IFNB* enhancers by IFN-γ priming (IFN-γ did not induce transcription of these genes). This aligns with prior research indicating that IFN-γ priming can induce epigenetic changes at the regulatory regions of inflammatory genes without initiating robust gene transcription ^20, 21, 22^. Curious about whether IFN-γ priming alone could increase chromatin accessibility at synergistic genes’ regulatory regions, we conducted upset plot analysis and found that 170 out of 517 (33%) of the synergy genes showed increased accessibility without corresponding changes in gene expression patterns, suggesting that IFN-γ priming alone can induce epigenetic changes on signature priming genes without activating transcription (Figure 3 E). As expected, the chromatin regions regulated by IFN- γ were characterized by an enrichment of ISRE/IRF1 (Homer *de novo* motif analysis) motifs (Figure 3 F). Interestingly, we also found significant enrichment for NF-κB motifs, further supporting the notion that IFN-γ priming induced chromatin remodeling of NF-κB target genes without activating inflammatory gene transcription (Figure 3 F).

**Figure 3:**
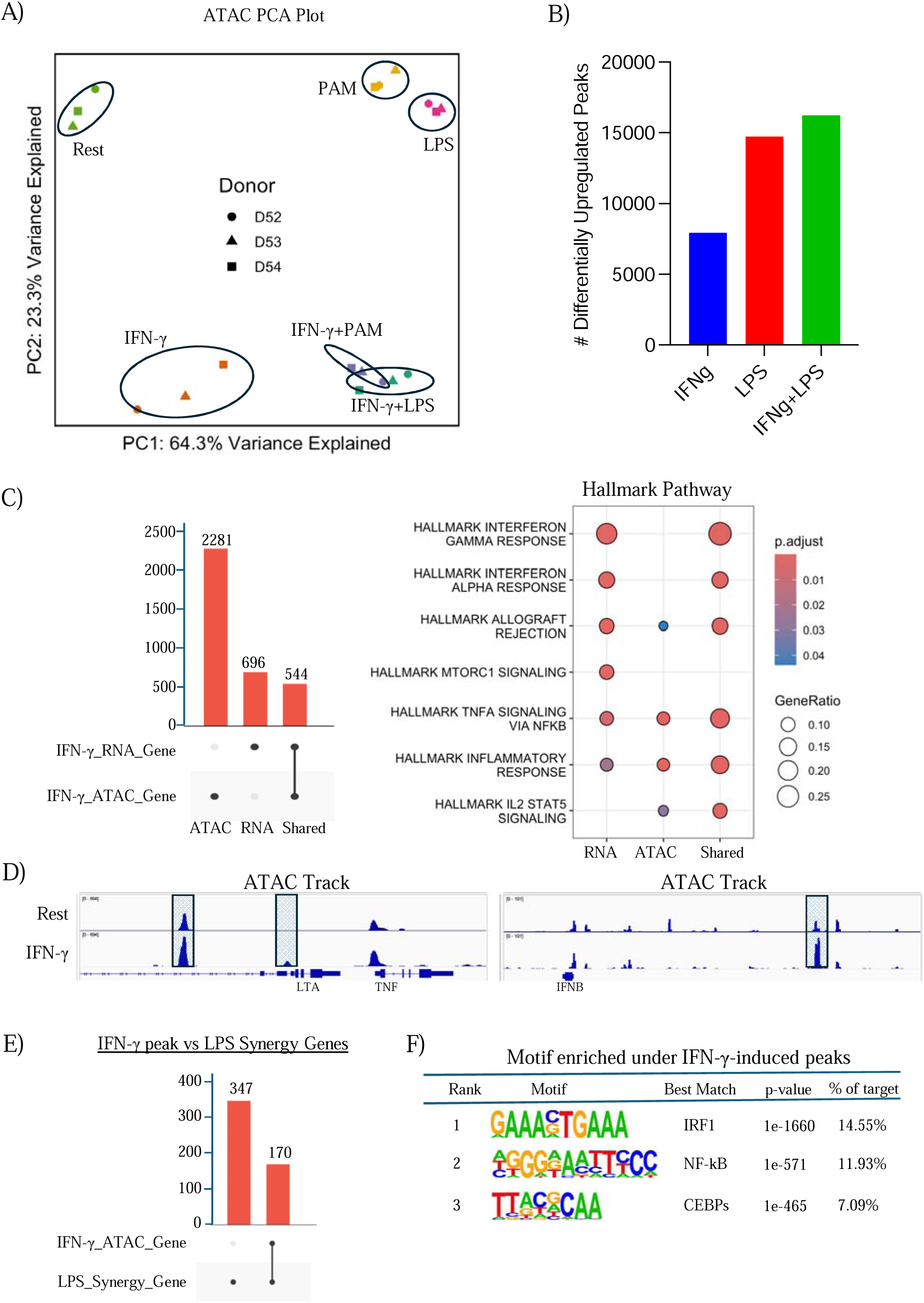
IFN-γ priming increased chromatin accessibility at a subset of signature priming genes regulatory elements. A) PCA plot of ATAC-seq data of IFN-γ-primed primary human monocytes challenged with indicated secondary stimuli. B) Bar plot depicting number of differentially upregulated ATAC peaks in IFN-γ, LPS and IFN-γ + LPS condition. (FDR ≤ 0.05 and 2-fold used as cut-off). C) UpSet plot showing overlap between genes associated with IFN-γ-induced ATAC peaks and IFN-γ-induced genes by RNA-seq. D) Representative IGV gene tracks of *de novo* IFN-γ-induced ATAC peaks at the regulatory region of genes displayed. No gene transcription detected. E) UpSet plot showing overlap between genes associated with IFN-γ-induced ATAC peaks and synergistically activated gene in IFN-γ-primed LPS-challenged monocytes as identified in RNA-seq analysis. F) *De novo* motif analysis results using HOMER of IFN- γ-induced ATAC.

We expanded our analysis to investigate whether IFN-γ priming enhanced LPS-induced ATAC peaks, similar to what was observed at gene level in the RNA-seq data. LPS significantly increased chromatin accessibility at 14,736 genomic regions (FDR ≤ 0.05 and >2-fold), while IFN- γ + LPS increased accessibility at 16,249 genomic regions (Figure 4A). Using a Venn diagram comparing LPS-induced peaks to IFN-γ + LPS-induced peaks, we found that out of the 16,249 IFN-γ + LPS-induced peaks, 5,279 (32.4%) were not identified as significant peaks in the LPS alone condition (termed Group i or ‘IFN-γ + LPS unique’, corresponding to *de novo* IFN-γ + LPS peaks) (Figure 4 A).

**Figure 4:**
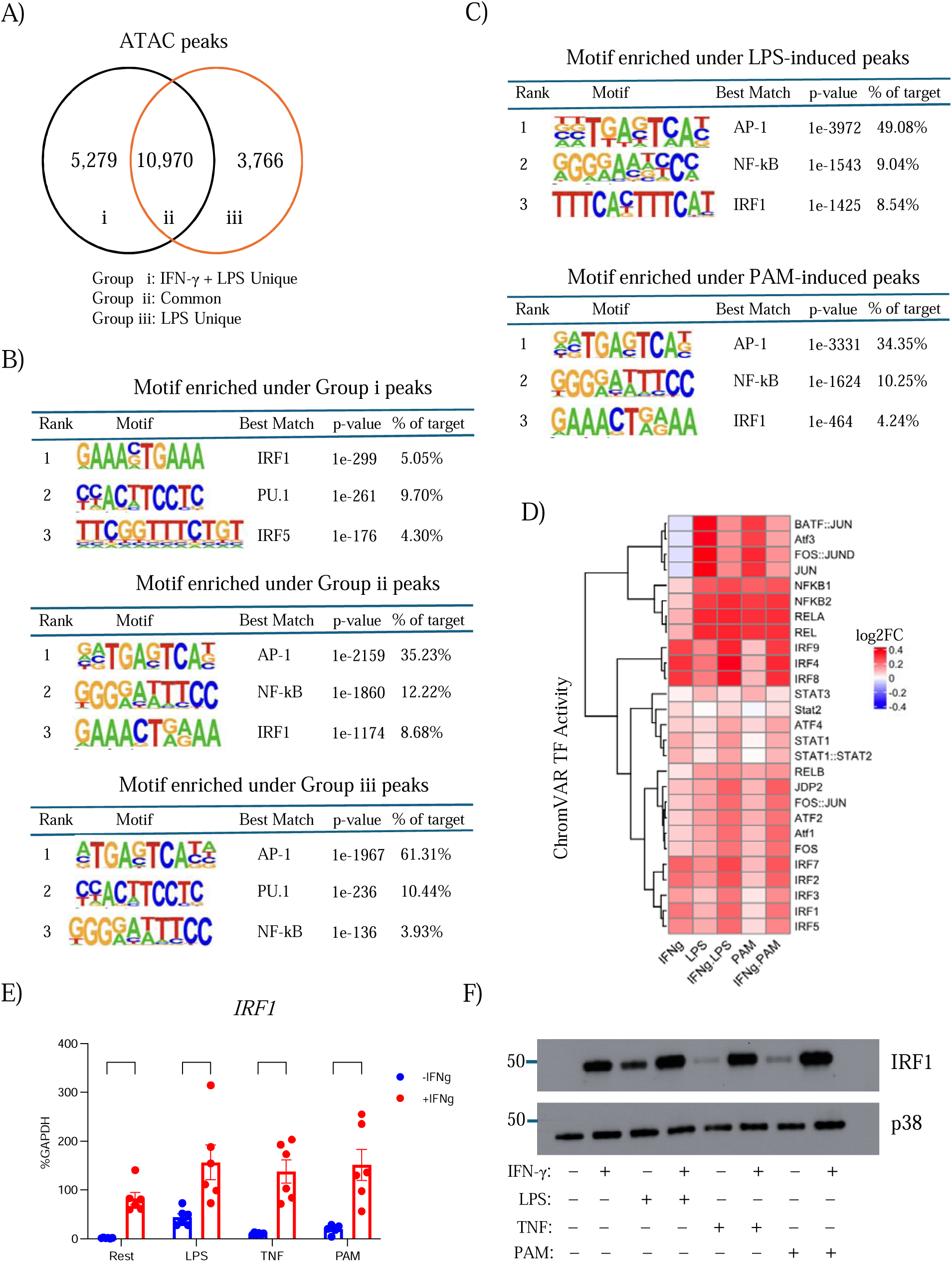
ATACseq analysis implicates IRFs in the IFN-γ-primed TLR response. A) Venn diagram illustrating numbers of LPS-induced ATAC peaks that are either called as significant or as non-significant by IFN-γ priming. B) *De novo* motif analysis results using HOMER of Groups identified in Figure. 2.5.A C) *De novo* motif analysis results using HOMER of ATAC peaks that are induced by LPS, PAM or TNF relative to resting. D) Heatmap of the differential TF activity scores derived from ChromVAR motif enrichment analysis of ATAC data for IFN-γ, LPS, IFN-γ +LPS, TNF, IFN-γ+TNF, PAM and IFN- γ+PAM treated human monocytes, compared to resting control. E) mRNA of *IRF1* genes was measured by qPCR and normalized relative to *GAPDH* mRNA in cells stimulated as indicated (dots correspond to independent donors; n = 6). Data are depicted as mean ± SEM. ** *p* < 0.001; **** *p* < 0.0001 by Two-way ANOVA with Tukey’s multiple comparisons test. F) Immunoblot of IRF1, and p38 using whole cell lysates of cells stimulated as indicated (representative blot from one out of 3 independent donors, p38 used as loading control)

Additionally, a substantial number of LPS-induced peaks (10,970 or 54.8%) were also present in the IFN-γ + LPS condition (termed Group ii or ‘Shared peaks’), whereas 3766 peaks (18.5%) were LPS induced peaks but not called as significant peaks in IFN-γ + LPS condition (Figure 4 A). Next, we used HOMER to perform *de novo* motif analysis to identify TF binding sequences enriched under ATAC peaks for each group (Figure 4 B). We found that both Group i and Group ii were enriched for IRF1 motifs, while Group ii and Group iii were enriched for AP-1 motifs. Group ii peaks, which are present in both LPS and IFN-γ + LPS conditions, were enriched for NF-κB motifs and also shared IRF motifs with Group i and AP-1 motifs with Group iii. These findings suggest that IFN-γ priming selectively suppresses a subset of LPS-induced ATAC peaks enriched for AP-1 motifs, enhances peaks enriched for a different subset of AP-1 and NF-κB motifs (Figure 4 B) and shows that peaks unique to the IFN-γ + LPS) condition are highly enriched in IRF motifs.

Similar to LPS, ATAC-seq analysis of IFN-γ-primed PAM-challenged monocytes mirrored the observations made with LPS challenge in IFN-γ-primed monocytes (data not shown). This suggests that TLR ligands utilizing similar adaptor proteins in their signaling cascades share common transcription factor motifs to induce synergistic gene expression. Additionally, we performed motif analysis using ChromVAR, a computational analysis program that generates TF activity scores. ChromVAR analysis confirmed HOMER results, showing that IFN-γ induced IRF activity, suppressed AP-1 activity and did not substantially affect TLR-induced NF-κB activity (Figure 4D) and that TLRs challenge alone highly induced IRF1 TF activity relative to the resting control (Figure 4 D). Both HOMER and ChromVAR results suggest that PAM and LPS can recruit IRF1 independently of IFN-γ signaling. Next, we wished to understand how co-treatment modulates IRF1 mRNA or protein levels compared to TLR-challenged resting monocytes. To test this, we measured IRF1 mRNA levels by RT-qPCR and found that IFN-γ was able to induce IRF1 expression at much higher level compared to TLRs challenge alone, but the induction was dramatically stronger in IFN-γ + LPS, and IFN-γ + PAM compared to TLR-challenged monocytes (Figure 4 E). Additionally, western blot analysis confirmed that TLR challenge alone induced relatively lower IRF1 protein levels at the whole cell protein level compared to IFN-γ-primed TLR- challenged monocytes (Figure 4 F). In accord with ChomVAR results, IRF1 expression and protein level were comparable in IFN-γ-primed monocytes when challenged LPS, or PAM (Figure 4 E-F). The results suggest a potential role for IRF1 and possibly other IRFs in the IFN-γ-primed inflammatory response.

### Transcriptional memory in IFN-γ-primed monocytes treated with iJAK baricitinib

We then sought to understand whether IRF1 played a role in the synergistic induction of inflammatory NF-κB target genes. Given that IRF1 expression levels were similar across dual-treated groups at both mRNA and protein levels and that IFN-γ alone induces significant levels of IRF1 (Figure 4 E-F), it is challenging to isolate IRF1’s potential contribution in inducing synergistic gene expression. To uncouple the contributions of IRF1 induced by IFN-γ and secondary challenges (TLR ligand), we inhibited IFN-γ signaling before the secondary challenges (Figure 5 A). Using baricitinib, an FDA-approved JAK1/2 inhibitor (iJAK), we found that IFN-γ- induced STAT1 activity was abrogated (assessed by phospho-STAT1 level) (not shown). Blocking IFN-γ signaling with baricitinib resulted in a near-complete loss of IFN-γ-induced IRF1 mRNA and protein levels (lane 3 vs lane 4) (Figure 5 B-C). IRF1 induced by LPS (lane 5 vs lane 6) was weakly suppressed by iJAK in unprimed monocytes (Figure 5 B). Strikingly, in IFN-γ-primed, iJAK-treated monocytes, IRF1 expression and protein levels were mostly maintained when challenged with LPS (Figure 5 B-C). This suggests that IRF1 mRNA and protein levels observed in the IFN-γ condition are strongly suppressed by baricitinib. In contrast, in the IFN-γ + LPS condition, the administration of iJAK had a less profound effect, and IRF1 protein levels in iJAK- treated (IFN-γ + LPS) cells were higher than in LPS-treated cells even in the absence of JAK inhibition. The results suggest that in primed cells, LPS-induced IRF1 can compensate for any loss of IFN-γ-induced IRF1 in the presence of iJAK (Figure 5 B-C). This finding aligns with the literature ^20, 21, 22^, which suggests that IFN-γ- induced IRF1 may serve a bookmarking mechanism to stably alter chromatin and enable TLR-induced IRF1 to bind and drive the induction of non-ISGs, such as NF-κB target genes. This suggests a role for IRF1 in augmenting monocytes for a robust inflammatory response upon subsequent TLR stimulation.

**Figure 5:**
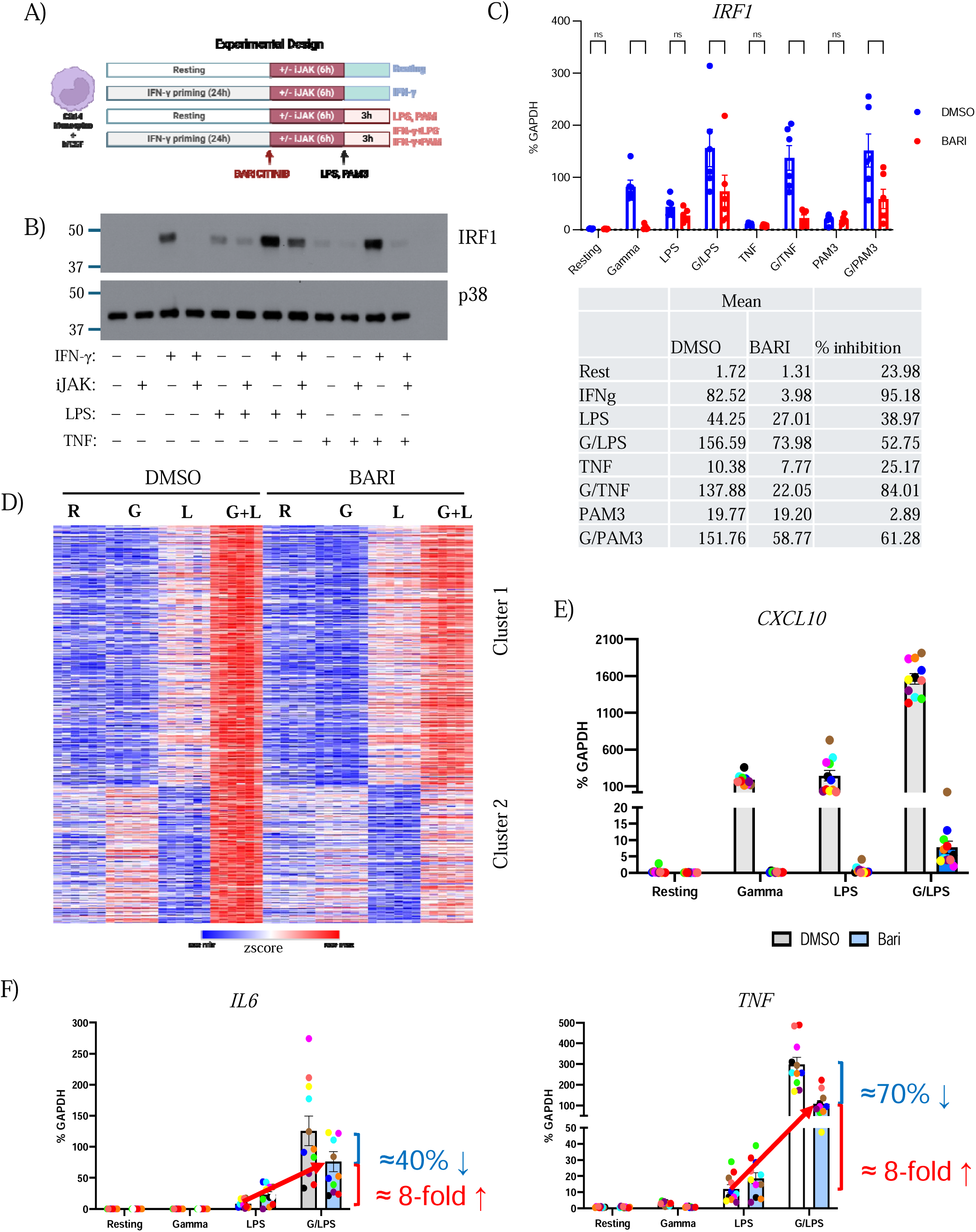
Transcriptional memory in IFN-γ-primed monocytes treated with iJAK baricitinib. A) Experimental design. Human monocytes were cultured –/+ IFN-γ (100 U/mL) for 24 hours, and cells were either treated with DMSO or Baricitinib for 6hours prior to stimulating with various inflammatory ligands for 3 hours. B) Immunoblot of IRF1, and p38 using whole cell lysates of cells stimulated as indicated (representative blot from one out of 3 independent donors, p38 used as loading control) C) mRNA of *IRF1* genes was measured by qPCR and normalized relative to *GAPDH* mRNA in cells stimulated as indicated (dots correspond to independent donors; n = 6). Data are depicted as mean ± SEM. **** *p* < 0.0001 by Two-way ANOVA with Tukey’s multiple comparisons test. D) Heatmap of synergistically activated gene in LPS model clustered by LPS vs IFN induced genes with or without Baricitinib treatment. E) mRNA of *CXCL10* genes was measured by qPCR and normalized relative to *GAPDH* mRNA in cells stimulated as indicated (dots correspond to independent donors; n = 11). Data are depicted as mean ± SEM. F) mRNA of *IL6* and *TNF* genes was measured by qPCR and normalized relative to *GAPDH* mRNA in cells stimulated as indicated (dots correspond to independent donors; n = 11). Data are depicted as mean ± SEM.

We next investigated the effect of iJAK on previously identified synergistically activated genes to determine whether the loss of IFN-γ-STAT1 signaling resulted in decreased expression of these synergy genes. RNA-seq analysis including iJAK conditions showed that IFN-γ-inducible synergy genes were strongly suppressed by iJAK (Fig. 5D, cluster 2), whereas LPS-induced synergy genes were partially resistant to iJAK (Figure 5 D, cluster 1). Interestingly, iJAK-resistant clusters harbored signature NF-κB target genes involved in the pathogenesis of inflammatory diseases such as rheumatoid arthritis (RA) (not shown).

We validated this finding by performing RT-qPCR and found that iJAK abrogated the expression of ISGs such as *CXCL10*, while LPS-induced NF-κB target genes such as *IL6* and *TNF* were resistant to JAK inhibition (Figure 5 E-F). Strikingly, superactivation of *IL6* and *TNF* in the IFN-γ + LPS condition was preserved when JAKs were inhibited (Figure 5F, compare 8-fold increase in column 8 to LPS alone in column 5). This persistence of high-level induction of inflammatory NF-κB target genes even after IFN-γ signaling is abrogated demonstrates ‘transcriptional memory’ and is consistent with a training effect that persists after the initial stimulus is terminated.

### Role of IRF1 in IFN-γ-mediated priming of inflammatory genes

We reasoned that the transcriptional memory, could be mediated by increased transcription factor (IRF, NF-κB, CEBPs, and others) activity ^21, 22^, and associated stable changes in chromatin, which are maintained even after IFN-γ signaling is abrogated. To test this notion, we extended our ATAC-seq analysis to include iJAK-treated conditions. As expected, IRF1 transcription factor (TF) activity driven by single treatments (IFN-γ or LPS) was diminished when cells were subjected to JAK inhibition (iJAK) (Figure 6A). Interestingly, IRF1 TF activity remained relatively unimpaired in the IFN-γ + LPS condition even with iJAK treatment, suggesting that IRF1 might contribute to the partial maintenance of inflammatory NF-κB target genes in the IFN-γ + LPS condition despite JAK inhibition (Figure 6A). We observed similar results in IFN-γ-primed PAM challenged monocytes (Figure 6B).

**Figure 6:**
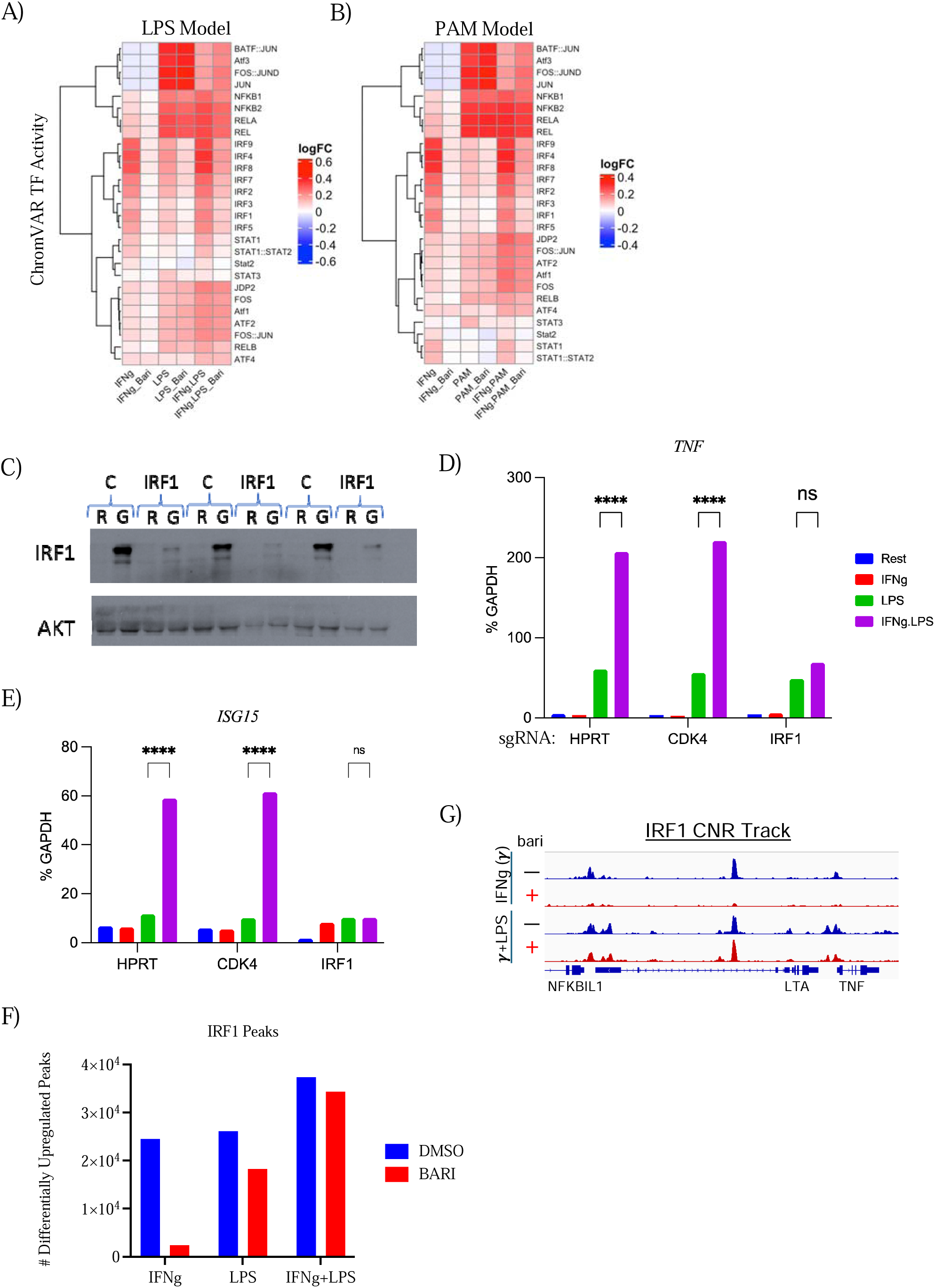
Role of IRF1 in IFN-γ-mediated priming of inflammatory genes. A) Heatmap of the differential TF activity scores derived from ChromVAR motif enrichment analysis of ATAC data for LPS model with Baricitinib treatment. B) Heatmap of the differential TF activity scores derived from ChromVAR motif enrichment analysis of ATAC data for PAM model with Baricitinib treatment. C) Immunoblot of IRF1 and AKT using whole cell lysates from *IRF1* or *HPRT*-edited cells (CRIPSR-Cas9 mediated edits) and stimulated with IFN-γ for 24h (n = 3 independent donors). D) mRNA of *TNF* was measured by qPCR and normalized relative to GAPDH mRNA after CRISPR-Cas9-mediated editing of IRF1, and stimulation as outlined in Figure 2.1.1.A (n=4 independent donors). E) mRNA of *ISG15* was measured by qPCR and normalized relative to GAPDH mRNA after CRISPR-Cas9-mediated editing of IRF1, and stimulation as outlined in Figure 2.1.1.A (n=4 independent donors). F) Bar plot depicting number of differentially upregulated IRF1 CUT&RUN peaks in IFN-γ, LPS and IFN-γ + LPS condition with and without Baricitinib treatment. (FDR ≤ 0.05 and 2-fold used as cut-off). G) Representative IGV gene tracks of IRF1 binding at the regulatory region of signature priming genes *TNF*.

To directly test the role of IRF1 in regulating synergy gene expression, we utilized CRISPR-Cas9 to edit the IRF1 gene locus in primary human monocytes, resulting in an almost complete loss of IRF1 protein expression (Figure 6C). We aimed to determine whether the loss of IRF1 resulted in a loss of synergistic activation of inflammatory NF-κB target genes such as *TNF*. Indeed, the loss of IRF1 completely abrogated the synergistic activation of *TNF* in the IFN-γ + LPS condition (Figure 6D). Similarly, we measured the mRNA level of the interferon-stimulated gene *ISG15* and found that the loss of IRF1 resulted in the loss of synergistic activation of ISG15 (Figure 6E). Overall, this provides strong genetic evidence for the role of IRF1 in the synergistic induction of *TNF* and *ISG15* in IFN-γ-primed monocytes when challenged with a TLR4 ligand.

This genetic evidence implicating IRF1 was corroborated by IRF1 CUT&RUN data showing that IRF1 binding peaks were highly sensitive to JAK inhibition under IFN-γ alone, but not under IFN-γ + LPS conditions (Figure 6 F and G). In accord with the idea that induction of synergy genes requires cooperation between different TFs in accessible chromatin ^21, 28, 29^, the results support cooperation between IRF1 and NF-κB at inflammatory gene loci.

## Discussion

We observed that IFN-γ priming significantly enhanced the expression of inflammatory genes such as *IL6*, *TNF, IL1B*, and interferon response genes like *CXCL10* when monocytes were stimulated with TLR ligands (TLR1/2, TLR4, TLR7/8). This synergistic activation suggests a broad regulatory effect of IFN-γ priming on the NF-κB pathway, extending beyond LPS-TLR4 interactions to include other TLR pathways. Our RNA-seq analysis further demonstrated that a substantial number of genes were synergistically activated across different TLR challenges, pointing to a common regulatory mechanism induced by IFN-γ priming.

Our ATAC-seq data provided insights into the epigenetic landscape of IFN-γ-primed monocytes. We found that IFN-γ priming increased chromatin accessibility at NF-κB target gene loci, suggesting that priming prepares these genes for increased transcriptional activation upon subsequent TLR stimulation. Differential motif analysis indicated significant enrichment of IRF1 and NF-κB motifs in accessible chromatin regions, implicating these transcription factors in the synergistic activation process, highlighting the importance of IRF1 in the transcriptional synergy observed with TLR ligands.

TLR signaling involves the MyD88-dependent and TRIF-dependent pathways. TRIF (TIR-domain-containing adapter-inducing interferon-β) is a key adaptor protein in TLR3 and TLR4 signaling. TLR3, which recognizes double-stranded RNA, exclusively signals through the TRIF-dependent pathway. Upon ligand binding, TLR3 recruits TRIF, activating downstream kinases such as TBK1 and IKKε, which phosphorylate IRF3. IRF3 then translocates to the nucleus to induce type I interferons (IFN-α and IFN-β) and other inflammatory cytokines. TLR4, which recognizes LPS, signals through both MyD88-dependent and TRIF-dependent pathways. Initially, TLR4 recruits MyD88, leading to early phase NF-κB and MAPK activation and pro-inflammatory cytokine production. Subsequently, TLR4 undergoes endocytosis and engages TRIF, leading to late-phase IRF3 and NF-κB activation and type I interferon production. In contrast, PAM exclusively activates the MyD88-dependent pathway. This results in NF-κB and MAPK activation and pro-inflammatory cytokine production but lacks type I interferon induction, unlike TLR3 and TLR4 via TRIF.

These differential pathways likely contribute to the observed differences in synergistic gene activation in IFN-γ-primed monocytes. LPS-induced synergy involves combined MyD88 and TRIF pathway activation, leading to robust and sustained NF-κB and IRF3/7 activation and synergistic induction of a broad set of pro-inflammatory genes, including type I interferons and IFN-stimulated genes. Conversely, PAM-induced synergy, by exclusively activating the MyD88- dependent pathway, may not achieve the same level of synergy in inducing type I interferons and IFN-stimulated genes but effectively induces pro-inflammatory cytokines through NF-κB activation. The interplay between MyD88 and TRIF pathways is crucial in determining the nature and extent of the inflammatory response, with LPS providing a broader and more sustained response compared to PAM. Understanding TRIF’s role and the differences between LPS and PAM signaling enhances understanding of the complex regulatory mechanisms governing the immune response in IFN-γ-primed monocytes.

Pathway analysis of synergistically activated genes revealed significant enrichment in inflammatory pathways, including TNF-NF-κB signaling and processes associated with rheumatoid arthritis. This enrichment underscores the relevance of our findings to inflammatory diseases and highlights potential therapeutic targets. The partial resistance of NF-κB target genes to JAK inhibition in the IFN-γ + LPS condition suggests that therapeutic strategies combining JAK inhibitors with other agents targeting NF-κB pathways could be more effective in managing inflammatory responses.

Our study raises several intriguing questions for future research. The exact mechanisms by which IFN-γ priming enhances chromatin accessibility and the roles of other transcription factors such as AP-1 and CEBPs in this process need further investigation. In conclusion, our findings provide a comprehensive view of the regulatory effects of IFN-γ priming on inflammatory gene activation in monocytes, revealing complex interactions between different signaling pathways and highlighting the critical role of chromatin remodeling in these processes. These insights contribute to a better understanding of immune regulation and offer potential avenues for therapeutic intervention in inflammatory conditions.

## Method Details

### Primary human monocytes

Deidentified buffy coats were purchased from the New York Blood Center following a protocol approved by the Hospital for Special Surgery Institutional Review Board. Peripheral blood mononuclear cells (PBMCs) were isolated using density gradient centrifugation with Lymphoprep (Accurate Chemical) and monocytes were purified with anti-CD14 magnetic beads from PBMCs immediately after isolation as recommended by the manufacturer (Miltenyi Biotec). Monocytes were cultured overnight at 37°C, 5% CO_2_ in RPMI-1640 medium (Invitrogen) supplemented with 10% heat-inactivated defined FBS (HyClone Fisher), penicillin-streptomycin (Invitrogen), L-glutamine (Invitrogen) and 20 ng/ml human M-CSF. Then, the cells were treated as described in the figure legends.

### Analysis of mRNA amounts (qPCR)

Total RNA was isolated using the RNeasy Mini Kit (QIAGEN, Cat#: 74106) following the manufacturer’s instructions. Reverse transcription of RNA into complementary DNA (cDNA) was performed using the RevertAid RT Reverse Transcription Kit (Thermo Fisher Scientific, Cat# K1691:) according to the manufacturer’s protocol, and the resulting cDNA was used for downstream analysis. For quantitative real-time PCR (qPCR), Fast SYBR Green Master Mix (Applied Biosystems, Cat#: 4385618) and a QuantStudio5 Real-time PCR system (Applied Biosystems) were used. CT values obtained from qPCR were normalized to the housekeeping gene *GAPDH*. Relative expression of target genes were calculated using the ΔCt method, where ΔCt represents the difference in threshold cycle values between the target gene and *GAPDH*. The results are presented as a percentage of *GAPDH* expression (100/2^ΔCt). Primer sequences used for the quantitative RT-qPCR reactions are provided in the Supplementary Table 1.

### Western blotting

For protein analysis we followed a previously reported method ^30^. Briefly, 2 x10^6^ human monocytes were washed with cold PBS after indicated treatments and harvested in 50uL cold lysis buffer containing Tris-HCl pH 7.4, NaCl, EDTA, Triton X-100, Na3VO4, phosSTOP EASYPACK, Pefabloc, and EDTA-free complete protease inhibitor cocktail. After a 10-minute ice incubation, cell debris was pelleted at 16,000xg at 4°C for 10 minutes. The soluble protein fraction was combined with 4× Laemmli Sample buffer (Bio-RAD, Cat#: 1610747) containing 2-mercaptoethanol and subjected to SDS-PAGE (electrophoresis) on 4–12% Bis-Tris gels. Following transfer of gels to polyvinylidene difluoride membranes, membranes were blocked in 5% (w/v) Bovine Serum Albumin in TBS with Tween-20 (TBST) at room temperature for at least one hour. Incubation with primary antibodies (diluted 1:1000 in blocking buffer) occurred overnight at 4°C. Membranes were washed three times with TBST and probed with anti-rabbit IgG secondary antibodies conjugated to horseradish peroxidase (diluted 1:2000 in blocking buffer) for one hour at room temperature. Enhanced chemiluminescent substrates (ECL western blotting reagents (PerkinElmer, cat: NEL105001EA) or SuperSignal West Femto Maximum Sensitivity Substrate (Thermo Fisher Scientific, Cat: 34095) were used for detection, followed by visualization on autoradiography film (Thomas Scientific, cat: E3018). For multi-protein detection on the same experimental filter while minimizing stripping and reprobing, membranes were horizontally cut based on molecular mass markers and the target proteins’ sizes. Restore PLUS western blotting stripping buffer (Thermo Fisher Scientific, Cat#: 46430) was applied for membranes requiring multiple primary antibody probes. Antibodies are listed in Supplementary Table 2.

### RNA sequencing

Libraries for sequencing were prepared using mRNA that was enriched from total RNA using NEBNext® Poly(A) mRNA Magnetic Isolation Module (New England Biolabs (NEB), Cat#: E7490L), and enriched mRNA was used as an input for the NEBNext Ultra II RNA Library Prep Kit (NEB, Cat#: E7770L), following the manufacturer’s instructions. Quality of all RNA and library preparations was evaluated with BioAnalyser 2100 (Agilent). Libraries were sequenced by the Genomic Resources Core Facility at Weill Cornell Medicine using a Novaseq SP flow cell, 50-bp pair-end reads to a depth of ∼20 - 40 million reads per sample. Read quality was assessed and adapters trimmed using FastQC and cutadapt. Reads were then mapped to the human genome (hg38) and reads in exons were counted against Gencode v38 with STAR Aligner. Differential gene expression analysis was performed in R using edgeR. Only genes with expression levels exceeding 4 counts per million reads in at least one group were used for downstream analysis. Benjamini-Hochberg false discovery rate (FDR) procedure was used to correct for multiple testing. Genes were categorized as upregulated if log2FC ≥ 1 and FDR ≤ 0.05 threshold was satisfied, downregulated if log2FC ≤ −1 and FDR ≤ 0.05. Heatmap with K-mean clustering was done using Morpheus web application and replotted using the R package pheatmap. Bioinformatic tools used are listed in Supplementary Table 3.

### Gene Set Enrichment Analysis (GSEA)

GSEA was conducted using the Broad Institute’s GSEA software (version 4.3.2). Log2 transformed counts per million (log2CPM) values of all expressed genes (CPM exceeding 4 in at least one group) in our RNA-seq dataset were used for gene ranking. The analysis utilized the h.all.v2023.2.Hs.symbols.gmt Hallmarks gene sets database. GSEA was performed with 1000 permutations, following default settings for the specific comparison outlined in the figure legend.

### Pathway analysis

The pathway analysis focused on investigating terms from Hallmark gene sets (https://www.gsea-msigdb.org/gsea/msigdb/human/collections.jsp#H) within a specific list of genes of interest. Gene sets representing pathways and biological processes were obtained using the R msigdbr package, which provided curated collections of genes associated with specific biological functions and pathways. Overrepresentation Analysis (ORA) was performed using the hypergeometric test to quantify the degree of enrichment, comparing the proportion of genes associated with a particular term within the list to the proportion in the entire genome. Implementation was conducted in R using the clusterProfiler ^31^ package, with subsequent visualization of results through dot plots or bar plots, effectively illustrating significantly enriched terms alongside their corresponding p-values.

### ATAC sequencing

The ATAC library preparation followed a previously reported method ^32^. Briefly, one million cells were lysed using cold lysis buffer (10 mM Tris-HCl, pH 7.4, 10 mM NaCl, 3 mM MgCl2, and 0.1% IGEPAL CA-630), and nuclei were immediately spun at 500x*g* for 10 min in a refrigerated centrifuge. The pellet obtained post nuclei preparation was resuspended in a transposase reaction mix consisting of 25 µl 2× TD buffer, 2.5 µl transposase (Illumina, Cat#: 20034198), and 22.5 µl nuclease-free water. The transposition reaction was carried out for 30 min at 37°C. Following transposition, the sample was purified using a MinElute PCR Purification kit. Library fragments were amplified using 1× NEB next PCR master mix and 1.25 M custom Nextera PCR primers as previously described ^33^, with subsequent purification using a Qiagen PCR cleanup kit, yielding a final library concentration of ∼30 nM in 20 µl. Libraries were amplified for a total of 10–13 cycles and subjected to high-throughput sequencing at the Genomic Resources Core Facility at Weill Cornell Medicine using the Illumina NovaSeq S1 Sequencer with 50-bp paired-end reads. Data from ATAC-seq experiments were derived from four independent experiments with different blood donors.

### ATAC-seq data analysis

For ATACseq data analysis, we utilized the TaRGET-II-ATACseq (https://github.com/Zhang-lab/TaRGET-II-ATACseq-pipeline) pipeline, available on a singularity image (ATAC_IAP_v1.1.simg), to process raw ATACseq data. The read alignments were performed against the GRCh38/hg38 reference human genome. Peak calling was conducted using MACS3 with the following parameters: “macs3 callpeak -f BAMPE -t replicate1 replicate2 replicate3 replicate4 -g hs -q 0.01 --keep-dup 1000 --nomodel --shift 0 --extsize 150”. A master consensus peak set was generated by merging the resulting peak files for each treatment condition, followed by merging peaks within 50bp of each other. Quantification of peaks to compare global ATACseq signal changes in the BAM files was conducted using the NCBI/BAMscale program. Raw count matrices were obtained utilizing the BAMscale program. Subsequent analysis utilized the HSS Genomic Core’s reproducible ATACseq analysis pipeline for peak filtering, annotation relative to genomic features, differential peak analysis, and enrichment of signal around specific motifs using ChromVAR.

### Motif Enrichment analysis (HOMER)

*De novo* transcription factor motif analysis was carried out using the motif finder program *findMotifsGenome* in the HOMER ^34^ package, focusing on the given peaks. Peak sequences were compared to random genomic fragments of the same size and normalized G+C content to identify enriched motifs in the targeted sequences.

### Interactive Genome Viewer (IGV)

To visualize the CUT and RUN, and ATAC-seq data, bigwig files were generated for each condition by merging replicate BAM files and then creating normalized coverage bigwig files relative to the sequencing depth using BAMscale ^35^. The normalized bigwig files were then visualized using the IGV browser from the Broad Institute ^36^.

### CUT and RUN

The Epicypher CUT and RUN kit (Cat#: 14-1048) was utilized following the manufacturer’s instructions. To create CUT and RUN libraries, fragmented DNA obtained from the CUT and RUN assay was processed using the NEBNext Ultra II DNA Library Prep Kit for Illumina (NEB, Cat# E7645L), following the manufacturer’s instructions. Libraries were pooled and sequenced (50bp paired end reads) to obtain at least 5 million reads/sample on the Illumina Novaseq SP or Novaseq S4 at the Genomic Resources Core Facility at Weill Cornell Medicine.

### CUT and RUN data analysis

A reproducible CUT and RUN analysis pipeline, CUT&RUNTools 2.0 ^37^ was used to process raw CUT and RUN data. In brief, the sequenced reads were aligned to the human genome (hg38) using bowtie2 ^38^. Peak calling was conducted using MACS2 with the following parameters: “macs2 callpeak -t replicate1 replicate2 replicate3 -g hs -f BAMPE -q 0.01--scale-to small --nomodel --keep-dup all”. The same approach used for ATAC-seq was applied for downstream analysis. Bioinformatic tools used are listed in Supplementary Table 3.

### CRISPR

For CRISPR-Cas9 genome editing in primary human monocytes, the Lipofectamine™ CRISPRMAX™ Cas9 Transfection Reagent (Invitrogen, Cat#: CMAX00008) was utilized. Initially, cells were resuspended at a concentration of 1.0 x 10^6 cells/mL in media with M-CSF (20ng/mL) and the Jak inhibitor baricitinib (1µM, to prevent activation of Jak-STAT signaling in response to transfected guide RNAs). The cells were then plated at a density of 100,000 cells per well in a 96-well plate and incubated at 37°C for 2-3 hours. Cas9 ribonucleoproteins (Cas9-RNPs) were generated using CRISPRMAX™ Cas9 Transfection Reagent, following the manufacturer’s protocols. Briefly, lyophilized single guide RNAs (sgRNAs) obtained from IDT-DNA (Supplemental Table 1) were reconstituted at a concentration of 50 µM in Nuclease-Free Duplex Buffer (IDT-DNA, Cat# 11-01-03-01). Subsequently, the two sgRNAs per gene were mixed with 61µM Cas9 protein (volume adjusted to achieve equal molar ratio) and incubated with opti-MEM and Cas9 Reagents (part of Lipofectamine kit) for 5 minutes at room temperature to form Cas9-RNPs at a final concentration of 5 µM. For delivery of Cas9-RNP to cells, Lipofectamine CRIPSRMAX lipid nanoparticles were diluted with Opti-mem and incubated for 1min at room temperature. Cas9-RNP and CRISPRMAX mix were incubated at a 1:1 ratio for 15min at room temperature. These Cas9-gRNA-Lipofectamine complexes were added immediately to the plated cells. The cells were then incubated overnight at 37°C. Following transfection, the media was changed to remove residual components, and the cells were further incubated for 24 hours before stimulating them with LPS for 3hours.

### Statistical Analysis

Graphpad Prism for Macs was used for all statistical analysis. Information about the specific tests used, and number of independent experiments is provided in the figure legends. Two-way ANOVA with Tukey’s correction for multiple comparisons was used for grouped data.

## Data and code availability

Sequencing data from this study will be deposited at GEO and will be publicly available from the date of publication. Any additional information required to reanalyze the data (including original code) reported in this paper is available from the first or last authors on request.

